# *In vivo* analysis of UNC-23 reveals residues critical for BAG2 domain function

**DOI:** 10.64898/2026.05.28.728440

**Authors:** Małgorzata Śledź, Ramakrishnan Ponath Sukumaran, Agata Szczepańska, Michał Turek

## Abstract

BAG2 is a co-chaperone that regulates Hsp70 activity and plays key roles in protein homeostasis and disease, yet the residue-level determinants of its stability and function remain poorly defined. Here, we use the *Caenorhabditis elegans* BAG2 homolog UNC-23 as an *in vivo* model to dissect these mechanisms. A forward genetic screen identified a missense mutation, UNC-23^C403Y^, that causes muscle deterioration without affecting developmental timing. Sequence and structural analyses revealed that C403 is highly evolutionarily conserved and located within a structurally conserved BAG domain. Furthermore, to systematically identify residues critical for protein stability, we combined evolutionary conservation with *in silico* stability predictions and tested selected substitutions *in vivo*. Despite strong conservation and predicted destabilization, most generated mutations were phenotypically silent, with only UNC-23^F416A^ producing a severe defect, revealing the limited predictive power of current approaches and a high tolerance of the BAG domain to perturbations. Comparative structural analysis further showed that, although the BAG domain architecture is conserved, its core stabilization relies on distinct interactions in invertebrates and vertebrates. Together, our results identify and refine residues contributing to UNC-23/BAG2 function, improving the current understanding of the BAG domain, and demonstrate that its architecture is maintained through distinct stabilizing interactions across evolution.

## Introduction

The maintenance of muscle integrity requires coordinated regulation of cytoskeletal organization, cell–matrix attachment, and protein homeostasis^1,2^. In *Caenorhabditis elegans*, body wall muscle cells are anchored to the hypodermis through specialized attachment structures that transmit contractile forces required for locomotion^3^. Disruption of these structures leads to progressive muscle detachment and degeneration, providing a genetically tractable system to study the molecular mechanisms that preserve tissue integrity^4,5^. While several structural components of muscle attachment have been characterized^6–8^, the contribution of protein quality control pathways to this process remains largely unexplored.

Protein homeostasis (proteostasis) is governed by molecular chaperones and their co-factors, which regulate protein folding, stabilization, and degradation^9^. Among these, the BAG (Bcl-2–associated athanogene) family of co-chaperones serves as a key modulator of Hsp70/Hsc70 activity, regulating protein folding and degradation pathways^10–12^. BAG2 is a member of this family characterized by a conserved BAG domain that mediates interaction with Hsp70 chaperones and regulates the fate of client proteins^13^. Through these interactions, BAG2 influences protein stability and degradation, thereby playing central roles in cellular stress responses.

Consistent with this function, BAG2 has been implicated in multiple biological contexts in which proteostasis is critical. One significant function of BAG2 is its role in modulating mutant p53 proteins, which are frequently associated with various cancers. Studies indicate that BAG2 promotes the accumulation of mutant p53 by inhibiting its degradation, thus enhancing its gain-of-function activities linked to tumorigenesis. This effect has been observed in different cancer types such as colorectal, lung, and breast cancers, where BAG2 expression levels correlate with tumour progression and poor prognosis^14–16^. BAG2 also acts as an anti-apoptotic factor, promoting cell survival under stressful conditions by preventing apoptosis^17^. Furthermore, BAG2 interacts with proteins involved in mitophagy, such as PINK1 and PARKIN, which are crucial for mitochondrial quality control. By inhibiting the ubiquitination of PINK1, BAG2 helps stabilize this protein, promoting its accumulation at the mitochondria, thereby facilitating the recruitment of PARKIN and enhancing the mitophagic clearance of damaged mitochondria^18^. BAG2 also has significant implications in neurodegenerative conditions, particularly concerning tauopathies. The ability of BAG2 to sequester tau and promote its degradation via ubiquitin-independent pathways reinforces its protective role against neurodegeneration^19^. However, despite its importance, the molecular determinants that govern BAG2 function, particularly the contribution of individual amino acid residues to its structural stability and interaction with chaperones, remain poorly understood.

The *C. elegans* protein UNC-23 is a conserved homolog of mammalian BAG2 and has been shown to play a critical role in maintaining muscle attachment and cellular integrity. Loss-of-function mutations in the *unc-23* gene result in detachment of muscle cells from the hypodermis, underscoring its importance in preserving muscle structure^5,20,21^. While these studies establish UNC-23 as an essential regulator of muscle integrity, the specific residues required for its activity and the extent to which its structural and functional properties are conserved with mammalian BAG2 remain unclear.

Here, we used an unbiased forward genetic approach to identify genes required for muscle integrity in *C. elegans*. This screen led to the identification of a novel missense mutation in *unc-23*, providing an entry point to dissect the function of this BAG2 homolog *in vivo*. Building on this discovery, we combined whole-genome sequencing, genome editing, evolutionary conservation analysis, and protein stability predictions to systematically identify residues critical for UNC-23/BAG2 function in muscle integrity. This approach revealed that not all evolutionarily conserved residues are required for this process, while uncovering conserved and divergent mechanisms that maintain the BAG domain across species.

## Results

### Generation of a novel *unc-23* mutant

To explore genetic factors affecting muscle integrity, we performed EMS mutagenesis on *C. elegans* worms expressing wrmScarlet fluorescent protein in body wall muscles (BWM). This mutagenesis approach aimed at generating a diverse set of mutations to study muscle function and integrity. Following screening, we identified a mutant displaying progressive muscle deterioration localized around the pharyngeal region (**Figure 1A-B**). To determine the underlying genetic basis of this phenotype, we conducted whole-genome sequencing (WGS) using the strategy described by Lehrbach et al.^22^. This approach involves outcrossing the mutant to a wild-type strain used for EMS mutagenesis and isolating multiple F2 progeny displaying the phenotype. To increase the reliability of our candidate gene identification, we performed two independent rounds of outcrossing. In each case, we selected F2 animals exhibiting the muscle deterioration phenotype and pooled their genomic DNA separately. Each pool was then subjected to whole-genome sequencing (**Figure 1C**).

**Figure 1.**
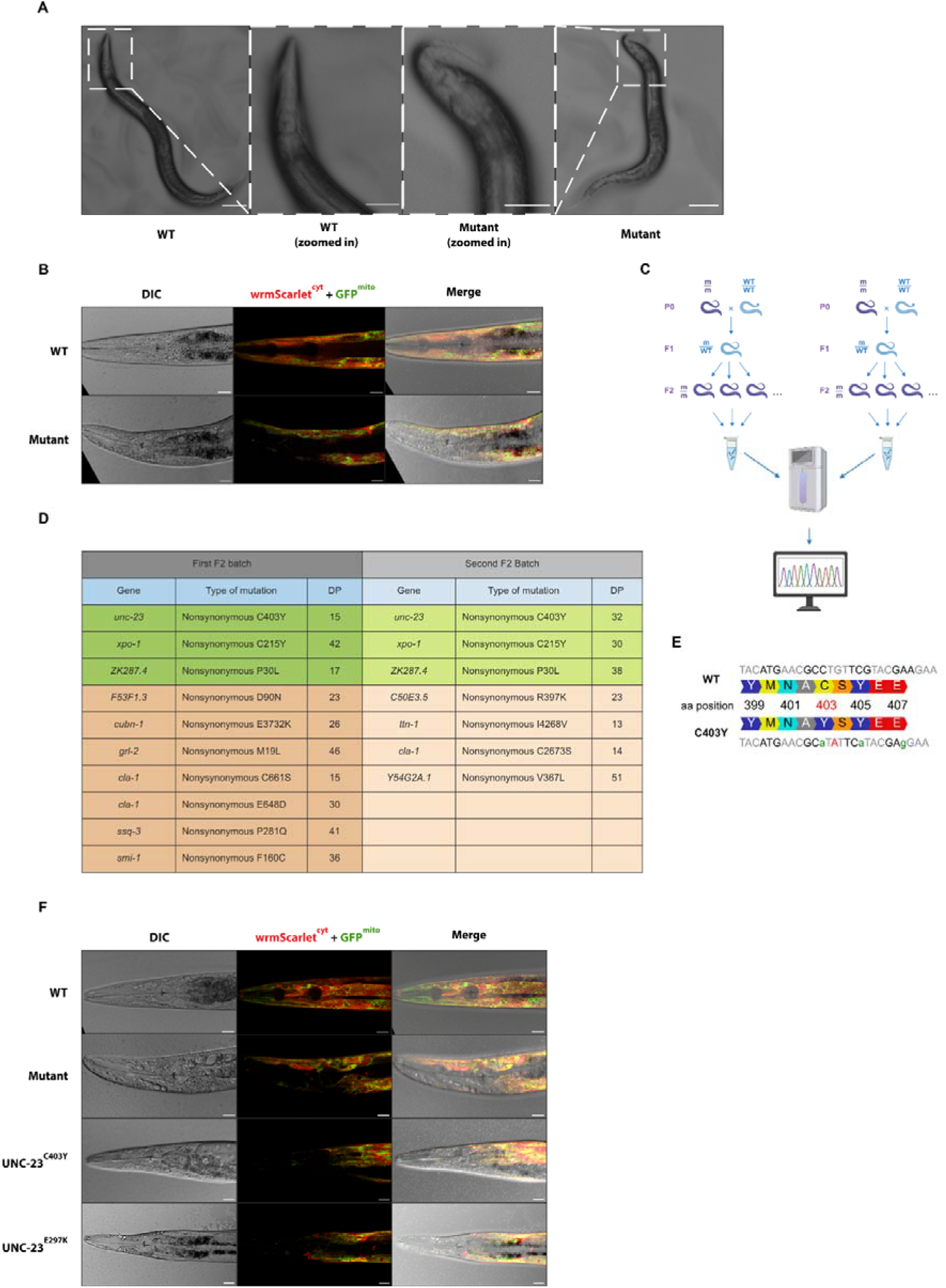
Identification of a causative C403Y mutation in *unc-23* leading to muscle deterioration. A Brightfield images of wild-type (WT) and mutant worms showing progressive muscle deterioration localized near the pharyngeal region. Insets show higher-magnification views of the affected area. Scale bars, 100 µm for the main images and 50 µm for the insets. B Representative DIC and fluorescence images of WT and mutant worms expressing cytosolic wrmScarlet and mitochondrial outer membrane-attached GFP in the body wall muscle. Merged images reveal disrupted muscle organization in the mutant. Scale bars are 20 µm. C Schematic of the whole-genome sequencing (WGS)-based strategy for mutation identification. Mutant worms were outcrossed to WT, and F2 progeny displaying the phenotype were selected, pooled, and sequenced in two independent experiments. Created with BioRender.com. D Whole-genome sequencing of two independent F2 pools identifies shared candidate mutations, including a nonsynonymous C403Y substitution in UNC-23. E CRISPR/Cas9-mediated generation of the *unc-23* C403Y allele, showing the nucleotide substitution responsible for the C403Y amino acid change and additional silent substitutions introduced in the crRNA target region to prevent Cas9 re-cutting. Red indicates the causative nucleotide substitution, whereas green indicates synonymous substitutions introduced for technical purposes that do not alter the amino acid sequence. F CRISPR/Cas9-engineered UNC-23^C403Y^ mutants exhibit muscle deterioration comparable to UNC-23^E297K^, confirming the causative role of the UNC-23^C403Y^ mutation. Scale bars are 20 µm.

By analysing the sequencing data from both independently outcrossed pools, we identified EMS-induced mutations by aligning sequencing reads to the reference genome and filtering for homozygous variants linked to the phenotype. Given that EMS typically induces hundreds of background mutations, we specifically searched for candidate mutations that were common to both backcrossed pools. This strategy allowed us to narrow down the list of putative causative mutations by eliminating those that were likely unrelated background mutations segregating differently across independent pools. Among the identified mutations, we found three genes with common variants in both pools: *unc-23* (nonsynonymous UNC-23^C403Y^), *xpo-1* (nonsynonymous XPO-1^C215Y^), and *ZK287.4* (nonsynonymous ZK287.4^P30L^) (**Figure 1D**). Based on literature evidence showing that mutations in the *unc-23* gene result in distorted muscle cell attachment to the hypodermis^21,23^, we prioritized a nonsynonymous missense mutation in *unc-23*, which led to a cysteine-to-tyrosine substitution at position 403 (UNC-23^C403Y^) in the UNC-23 protein, a homolog of the BAG2 protein. Given the established role of UNC-23 in maintaining cellular and muscle integrity^21^, we hypothesized that this mutation was responsible for the observed phenotype and proceeded with functional validation.

To confirm that this mutation was responsible for the observed muscle deterioration phenotype, we used CRISPR/Cas9 genome editing^24^ to precisely recreate the C403Y mutation in an otherwise wild-type background (**Figure 1E**). The resulting CRISPR-engineered mutants were verified by sequencing and exhibited the same progressive muscle deterioration phenotype as the EMS-induced mutant (**Figure 1F**), strongly supporting the conclusion that the C403Y mutation in UNC-23 is causative. Furthermore, we observed that the classical *unc-23* allele *e25* (nonsynonymous UNC-23^E297K^)^21,23^ exhibits a similar muscle deterioration phenotype, further supporting the notion that the C403Y mutation in UNC-23 leads to defects in muscle integrity (**Figure 1F**).

### The C403Y mutation impairs body and muscle morphology without affecting developmental timing

To assess the severity of the UNC-23^C403Y^ mutation, we quantified body and muscle morphology as well as developmental progression. We measured the width of the worm’s head and body wall muscles adjacent to the pharynx, from both the ventral and dorsal sides (**Figure 2A**), and compared these measurements with WT and UNC-23^E297K^ mutants. Our analysis revealed that the overall head and BWM width were significantly reduced in UNC-23^C403Y^ mutants compared to WT animals (**Figure 2B-D**). Interestingly, the head and dorsal BWM width in UNC-23^C403Y^ mutants were comparable to those in UNC-23^E297K^ animals, whereas the ventral BWM appeared slightly wider (**Figure 2B-D**).

**Figure 2.**
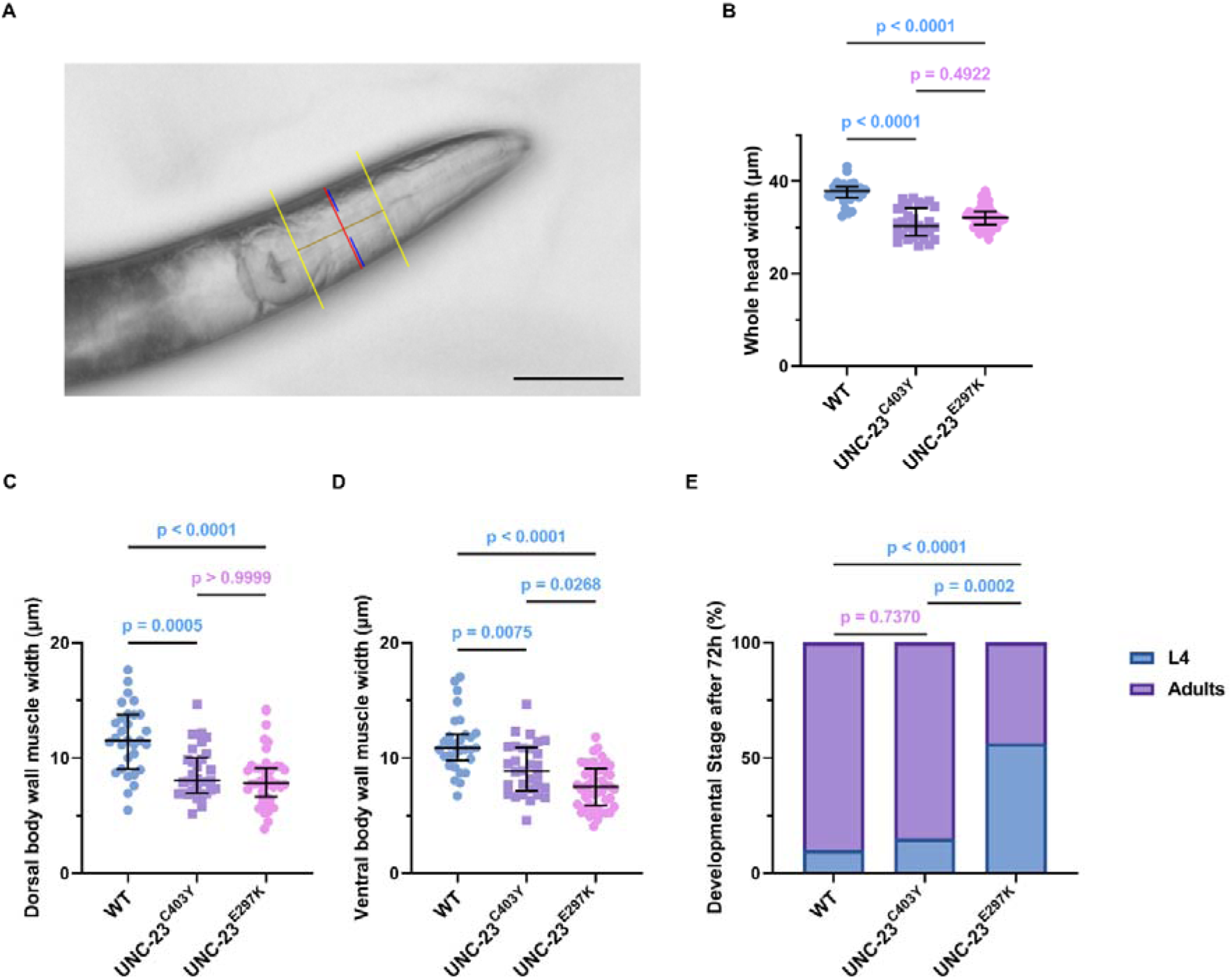
The UNC-23^C403Y^ mutation impairs body and muscle morphology without affecting developmental timing. A Representative image illustrating the measurement of head and body wall muscle (BWM) width adjacent to the pharynx. Measurements were taken from both dorsal and ventral sides. Yellow lines indicate guide lines, the red line marks head width, and the blue line marks muscle width. The ventral side was identified based on the position of the uterus. Scale bar, 50 μm. B Quantification of whole head width in wild-type (WT), UNC-23^C403Y^, and UNC-23^E297K^ animals. Head width is reduced in UNC-23^C403Y^ and UNC-23^E297K^ mutants compared to WT. (n = 30-45 worms, N = 3 independent experiments) C, D Quantification of dorsal (C) and ventral (D) BWM width. UNC-23^C403Y^ mutants exhibit reduced muscle width compared to WT, with values comparable to UNC-23^E297K^ mutants for dorsal measurements and slightly increased ventral width relative to UNC-23^E297K^. (n = 30-45 worms, N = 3 independent experiments) E Developmental progression assessed 72 h after synchronization. UNC-23^C403Y^ mutants develop at a rate comparable to WT, whereas UNC-23^E297K^ animals show delayed development. (n = 40-41 worms, N = 3 independent experiments) Data information: Data are presented as medians with interquartile ranges. Non-significant p-values (p > 0.05) are highlighted in pink, while significant p-values (p < 0.05) are shown in blue. Statistical significance was assessed using the Kruskal-Wallis test with Dunn’s multiple comparisons test (**B-D**) and Fisher’s exact test (**E**).

We next investigated whether the UNC-23^C403Y^ mutation affected developmental timing. To this end, we age-synchronized worms by hand-picking embryos at the 3-fold stage and assessed their developmental stage after 72 hours. The results showed that UNC-23^C403Y^ mutants developed at the same rate as WT worms, while UNC-23^E297K^ mutants exhibited a noticeable developmental delay compared to both WT and UNC-23^C403Y^ animals (**Figure 2E**).

### Cysteine 403 in UNC-23 is highly conserved across the animal kingdom

To assess whether cysteine 403 in UNC-23 is preserved across species, we performed a multiple sequence alignment of BAG2 homologs from a range of invertebrate and vertebrate organisms, including humans, using Clustal Omega^25^. The analysis showed that this cysteine residue is present in all examined homologs, indicating strong conservation and suggesting an important role in protein structure and function (**Figure 3A**). To complement this analysis, we also compared the AlphaFold-predicted structures of *C. elegans* UNC-23 and human BAG2. These models revealed that the overall fold is highly similar, with the BAG domain in both proteins adopting nearly identical three-dimensional structures (**Figure 3B**). To quantify the structural similarity between *C. elegans* UNC-23 and human BAG2, we performed a global structural alignment of their AlphaFold-predicted models using the Matchmaker tool in UCSF ChimeraX. Although the root-mean-square deviation (RMSD) across all 197 atom pairs was 8.919 Å (largely due to divergence in flexible termini and non-conserved regions) the core functional architecture was remarkably conserved. Focusing on a pruned subset of 99 residues encompassing the BAG domain and its immediate flanking regions, the RMSD dropped to just 0.733 Å. This sub-angstrom deviation confirms that the BAG domain adopts an almost identical three-dimensional fold in these evolutionarily distant species.

**Figure 3.**
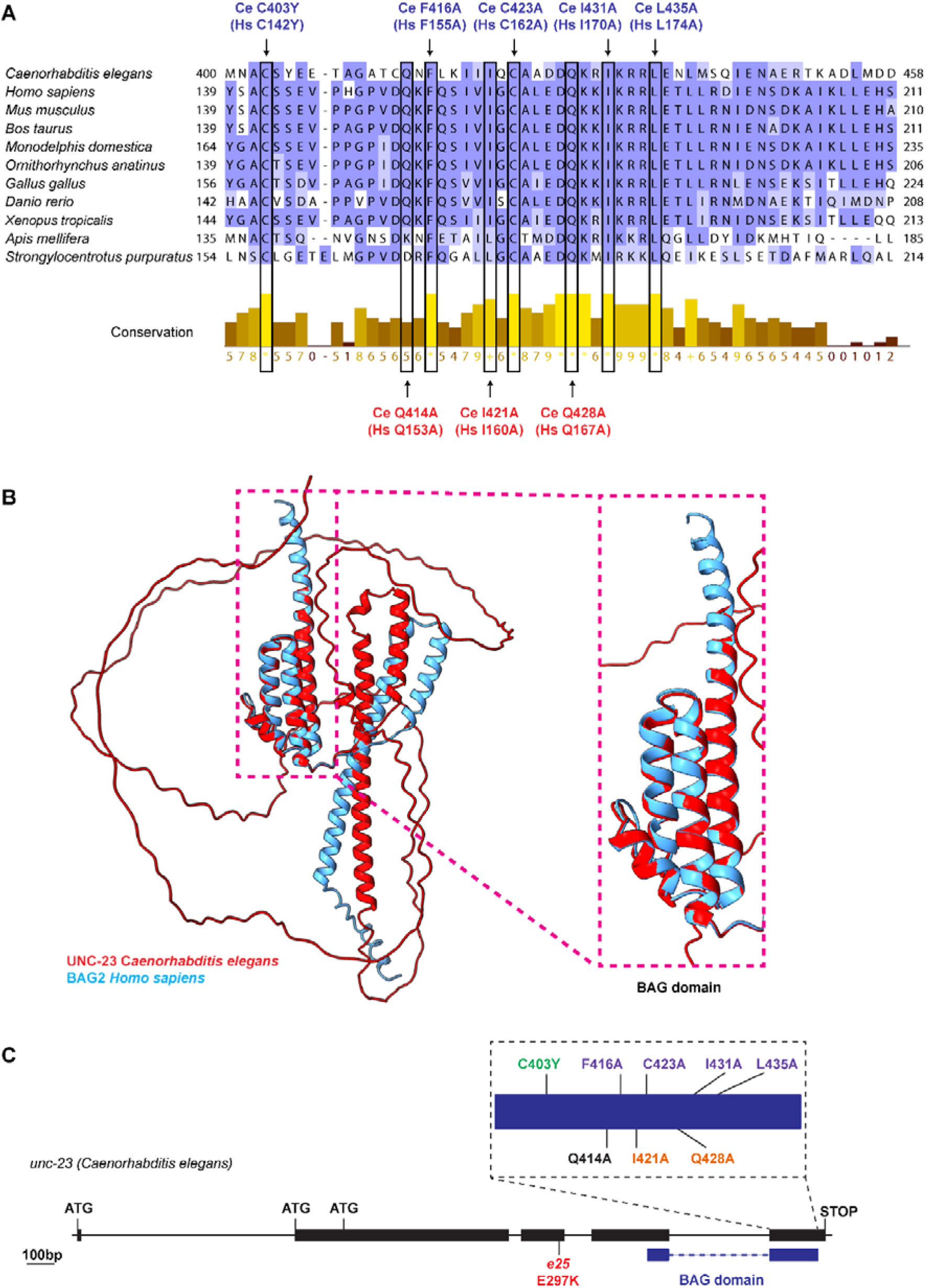
Cysteine 403 in *C. elegans* UNC-23 is highly conserved and located within a structurally conserved BAG domain of UNC-23/BAG2. A Multiple sequence alignment of BAG2 homologs from invertebrate and vertebrate species shows that cysteine 403 in *C. elegans* UNC-23 (corresponding to C142 in human BAG2) is highly conserved, together with additional residues within the BAG domain analysed in this study. Residues newly investigated here are marked in blue, whereas residues previously examined by Xu et al.^13^ are marked in red. For *C. elegans* (Ce) residues, the corresponding human (Hs) BAG2 residues are indicated in parentheses. B Structural alignment of AlphaFold-predicted *C. elegans* UNC-23 and human BAG2 proteins demonstrates strong conservation of the overall fold, with near-identical architecture of the BAG domain despite divergence in non-conserved regions. C Schematic representation of the *unc-23* gene showing the positions of residues analysed in this study. Black boxes indicate exons, connecting lines indicate introns, and the blue bar marks the BAG domain. The identified C403Y mutation is shown in green, the classical *unc-23* mutation is shown in red, previously studied mutations used here as positive controls are shown in orange, a previously studied mutation used as a negative control is shown in black, and newly generated mutations selected based on evolutionary conservation and predicted contribution to BAG domain stability are shown in purple. Scale bar is 100 bp.

The conservation of cysteine 403 supports its functional importance in UNC-23 and is consistent with the pronounced phenotype observed in UNC-23^C403Y^ mutants. This residue likely contributes to UNC-23 function, potentially through interactions with Hsc70 (HSP-1 in *C. elegans*)^13^, a molecular chaperone involved in protein folding and degradation^26^, and chaperone-mediated autophagy and the endosomal microautophagy^27^. BAG2 has been shown to modulate Hsc70 activity by preventing client protein ubiquitination and degradation, a process essential for maintaining proteostasis^13^. Notably, C403 in UNC-23 corresponds to C142 in human BAG2, for which structural evidence demonstrates participation in Hsc70 interaction^13^, suggesting that this conserved cysteine contributes to stabilizing this interaction, with its disruption leading to loss of protein function.

Disruption of BAG2 interaction with Hsc70 impairs its function and leads to defects in protein quality control and cellular stress responses. In addition to protein–protein interactions, protein stability itself has emerged as a critical determinant of function^28,29^. Destabilizing missense mutations can impair protein activity by disrupting structural integrity, often leading to pathological consequences, including cancer and neurodegenerative diseases. Despite the central role of BAG2 in proteostasis, residues specifically contributing to its structural stability remain poorly defined. A better understanding of how individual amino acids influence BAG2 stability and function is therefore essential for elucidating its molecular mechanisms.

Given the strong phenotype observed upon mutation of UNC-23 at key residues, we leveraged this model to identify conserved amino acids essential for BAG2 function in maintaining muscle integrity. By targeting these residues and assessing their impact on worm physiology, we aimed to gain deeper mechanistic insights into BAG2 function in humans. By evaluating mutations within a living organism, our strategy captures the multifaceted interactions and systemic effects that are often absent in isolated biochemical assays, providing a more comprehensive understanding of protein function and its implications for human health.

### Validation of the evolutionary conservation strategy through functional assays

To validate our approach, we assessed the reliability of our strategy by introducing mutations in both highly conserved and less conserved residues within the Hsc70 contact interface of UNC-23/BAG2. Highly conserved residues, previously demonstrated to be functionally essential, were mutated and served as positive controls, whereas a less conserved residue, shown to have minimal impact on function, was used as a negative control^13^. Specifically, as positive controls, we introduced mutations in *C. elegans* UNC-23^I421A^ (corresponding to BAG2^I160A^ in humans) and UNC-23^Q428A^ (corresponding to BAG2^Q167A^ in humans) (**Figure 3A and C**), both of which are critical residues forming the Hsc70 contact interface with the UNC-23/BAG2 BAG domain. As expected, these mutations resulted in a strong muscle deterioration phenotype, consistent with their known importance in the BAG domain–Hsc70 interface (**Figure 4**). Conversely, as a negative control, we introduced mutation UNC-23^Q414A^ (corresponding to BAG2^Q153A^ in humans) (**Figure 3A and C**), another residue located within the Hsc70 interface but previously shown to have minimal functional significance^13^. As anticipated, this mutation did not produce any detectable phenotype (**Figure 4**), further supporting the specificity of our approach. Taken together, these findings, along with the high structural similarity between *C. elegans* UNC-23 and human BAG2, particularly within the BAG domain (**Figure 3**), support the use of UNC-23 as a valuable model for investigating the structural and functional conservation of BAG2 in relation to its human homolog.

**Figure 4.**
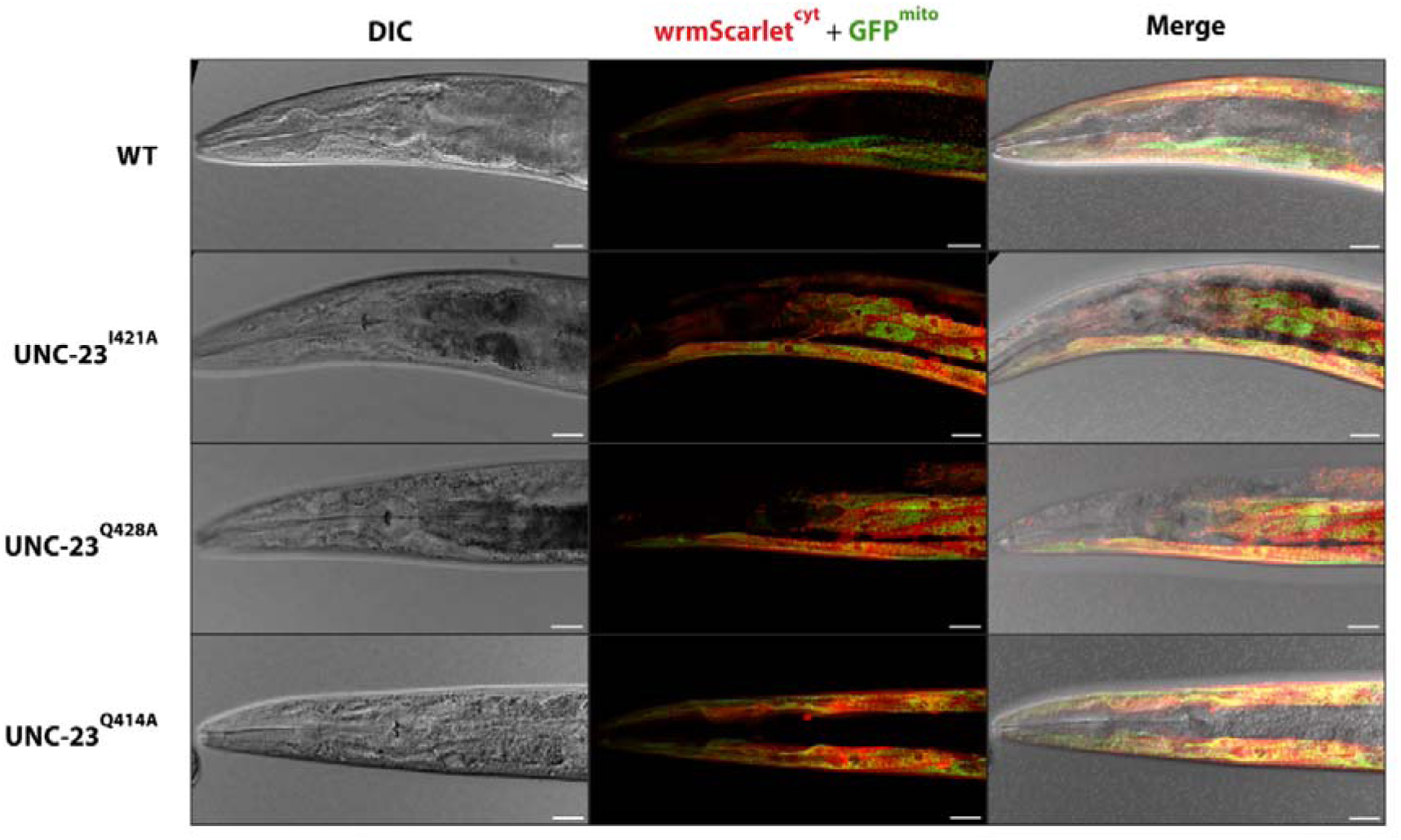
Mutations in conserved Hsc70-interacting residues of UNC-23 disrupt muscle integrity, whereas a less conserved residue has no detectable effect. Mutations in conserved residues of UNC-23, UNC-23^I421A^ and UNC-23^Q428A^, result in pronounced muscle deterioration, consistent with their essential role in BAG domain–Hsc70 interaction. In contrast, mutation of a less critical residue, UNC-23^Q414A^, does not produce detectable muscle defects. Representative DIC and fluorescence images of WT and mutant worms expressing cytosolic wrmScarlet and mitochondrial outer membrane-attached GFP in the body wall muscle. Merged images reveal disrupted muscle organization in mutants. Scale bars are 20 µm.

### Identification of residues predicted to contribute to UNC-23/BAG2 structural stability

In order to identify residues that are crucial for maintaining protein structure and are likely not involved in interactions with molecular partners, we utilized DDMut analysis, which predicts the impact of amino acid substitutions on protein stability by calculating changes in free energy (ΔΔG) upon mutation^30^. Using this approach, we assessed the effect of alanine substitutions on *C. elegans* UNC-23 and human BAG2 proteins, allowing us to predict which residues are essential for preserving the structural integrity of BAG2.

The DDMut analysis revealed that the top 15 most stabilizing residues in both *C. elegans* UNC-23 and human BAG2 are predominantly hydrophobic amino acids, such as leucine, isoleucine, phenylalanine, and valine (**Supplementary Table 1**). This finding is consistent with previous studies demonstrating that hydrophobic residues play a crucial role in stabilizing the three-dimensional structure of proteins by forming a hydrophobic core that minimizes solvent exposure^31,32^. Disrupting these hydrophobic interactions often leads to significant destabilization and misfolding. In *C. elegans* UNC-23, the most destabilizing substitutions included UNC-23^F416A^ (−3.54 kcal/mol), UNC-23^L435A^ (−2.32 kcal/mol), and UNC-23^I420A^ (−2.15 kcal/mol), indicating that these residues are predicted to make major contributions to structural stability. Similarly, in human BAG2, the most destabilizing mutations involved BAG2^L174A^ (−2.88 kcal/mol), BAG2^F155A^ (−2.51 kcal/mol), and BAG2^I170A^ (−2.09 kcal/mol). Examples of changes in interaction due to alanine replacement for F416 *C. elegans* UNC-23 and corresponding F155 in the human BAG2 homolog are shown in **Figure 5A and B**, illustrating the impact of these substitutions and their potential consequences on protein stability and function. The overlap in predicted destabilizing mutations between UNC-23 and BAG2 (**Supplementary Table 1**) indicates that core features governing BAG domain stability are conserved, underscoring the value of *C. elegans* as a model for investigating BAG2 function *in vivo*.

**Figure 5.**
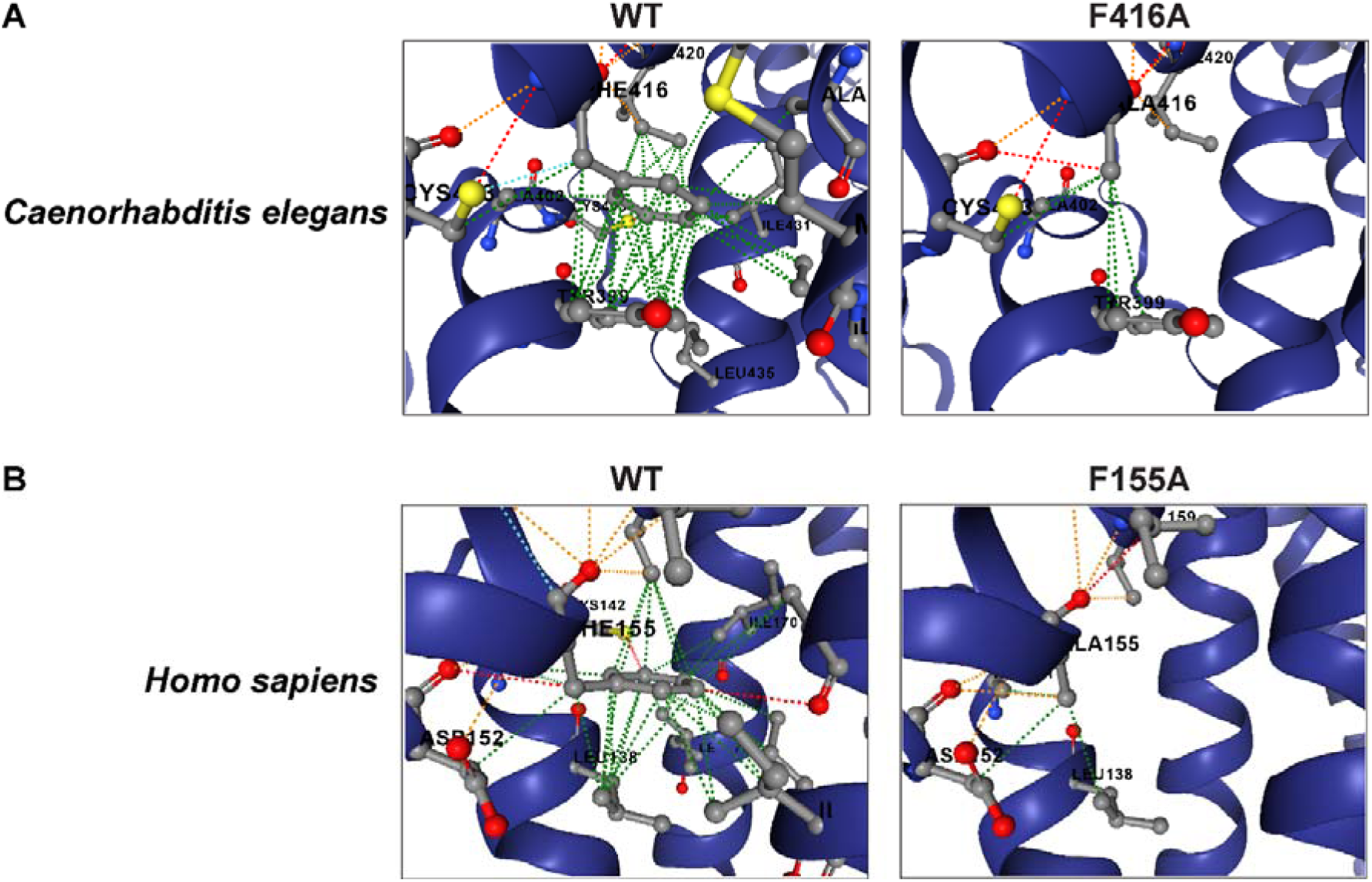
Hydrophobic core residues are predicted to be critical for UNC-23/BAG2 structural stability. A, B Alanine substitutions of conserved hydrophobic residues, (A) UNC-23^F416A^ and its (B) human BAG2 counterpart BAG2^F155A^, are predicted to disrupt stabilizing interactions within the protein core. Structural representations illustrate loss of key hydrophobic contacts upon mutation, consistent with strong destabilizing effects predicted by DDMut analysis.

To further refine our analysis, we cross-referenced data from evolutionary conservation and protein stability predictions to identify residues that are identical from invertebrates to mammals and ranked among the top 15 most crucial residues for protein stability (**Supplementary Table 1**). Based on this approach, we selected four out of five top highly conserved amino acids that met both criteria and introduced alanine substitutions in *C. elegans* UNC-23 to assess their functional impact. These residues were UNC-23^F416A^ (corresponding to BAG2^F155A^ in humans), UNC-23^C423A^ (corresponding to BAG2^C162A^ in humans), UNC-23^I431A^ (corresponding to BAG2^I170A^ in humans), and UNC-23^L435A^ (corresponding to BAG2^L174A^ in humans) (**Figure 3A and C**).

Unexpectedly, only the UNC-23^F416A^ substitution resulted in a pronounced muscle deterioration phenotype, while the other three mutations did not produce detectable effects (**Figure 6**). Given the strong evolutionary conservation of cysteine 423, isoleucine 431, and leucine 435 from sea urchins to humans, these residues are likely to be functionally relevant. However, it is possible that the substitutions introduced here cause only subtle destabilizing effects that are insufficient to compromise muscle integrity under the conditions tested. It also remains possible that these mutations affect other aspects of UNC-23 function not revealed by our current assays.

**Figure 6.**
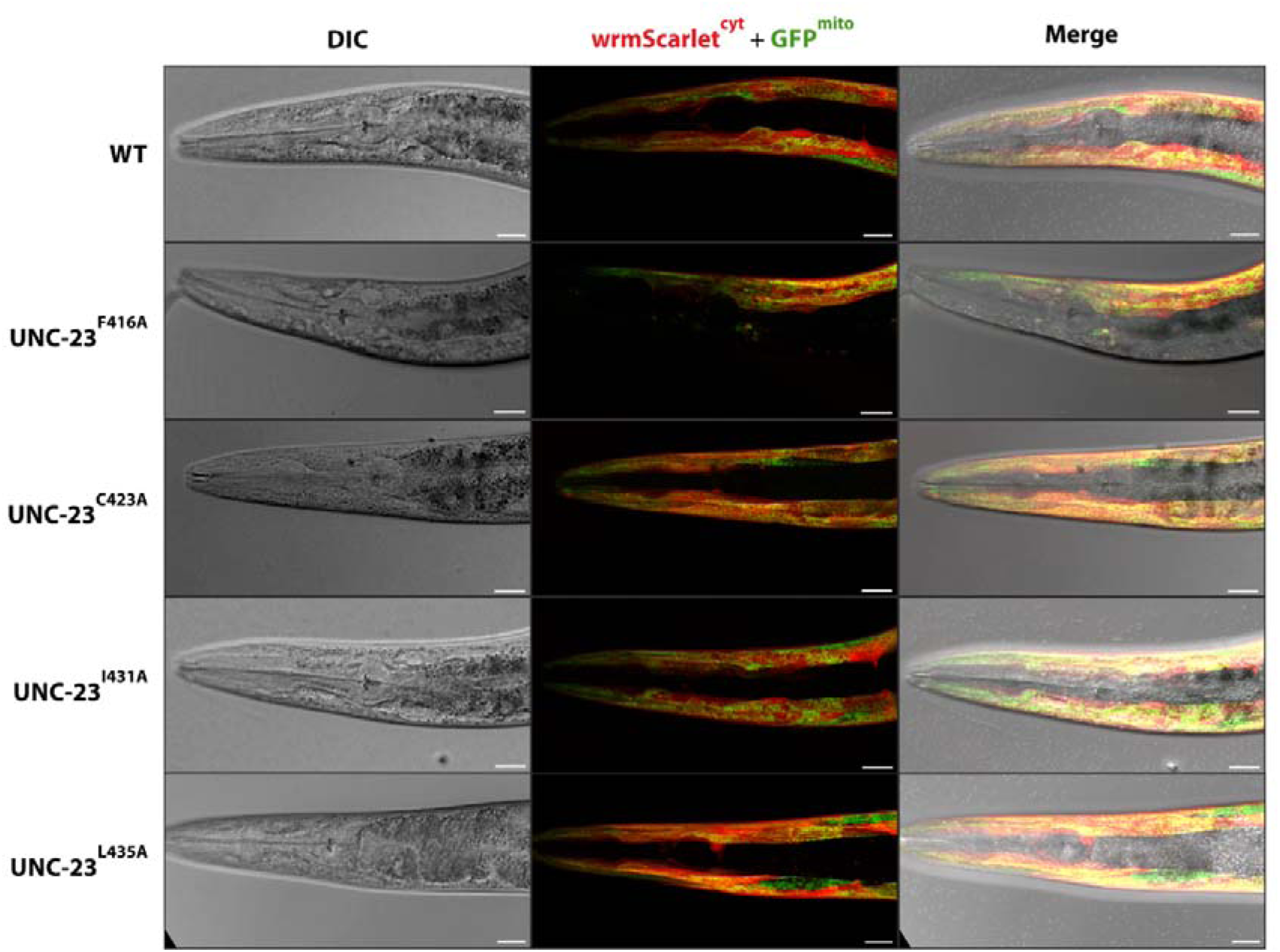
Only a subset of conserved residues predicted to stabilize UNC-23 is required for muscle integrity. Among conserved residues predicted to contribute to structural stability, mutation of UNC-23^F416A^ results in pronounced muscle deterioration, whereas mutations UNC-23^C423A^, UNC-23^I431A^, and UNC-23^L435A^ do not produce detectable muscle defects. Representative DIC and fluorescence images of WT and mutant worms expressing cytosolic wrmScarlet and mitochondrial outer membrane-attached GFP in the body wall muscle. Merged images reveal disrupted muscle organization in mutant UNC-23^F416A^. Scale bars are 20 µm.

### Conserved structural architecture of the BAG domain with divergent core stabilization mechanisms

Our analysis also demonstrates that while the overall structure of the BAG domain predicted by the AlphaFold is highly conserved between invertebrates and vertebrates (**Figure 3B**), and a large number of residues remain identical (**Figure 3A**), there is a key difference in the most crucial interactions stabilizing the core of the domain. Specifically, in invertebrates such as *C. elegans*, sea urchin, and honeybee, the core stabilization appears to involve a π–π stacking interaction between phenylalanine and tyrosine (F416 and Y399 in *C. elegans*) (**Figure 7**), or an equivalent interaction involving a phenylalanine in other invertebrate species. In contrast, in vertebrates, including humans, the stabilization occurs through hydrophobic interactions between phenylalanine (F155) and leucine (L174) rather than π–π stacking (**Figure 7**). Despite this variation in the specific stabilizing interaction, the overall structural arrangement of the BAG domain remains conserved. This finding suggests that while the mechanistic details of core stabilization have diverged, the functional integrity of the domain has been maintained through distinct but equivalent interactions, reflecting an evolutionary adaptation that preserves BAG2 function across species.

**Figure 7.**
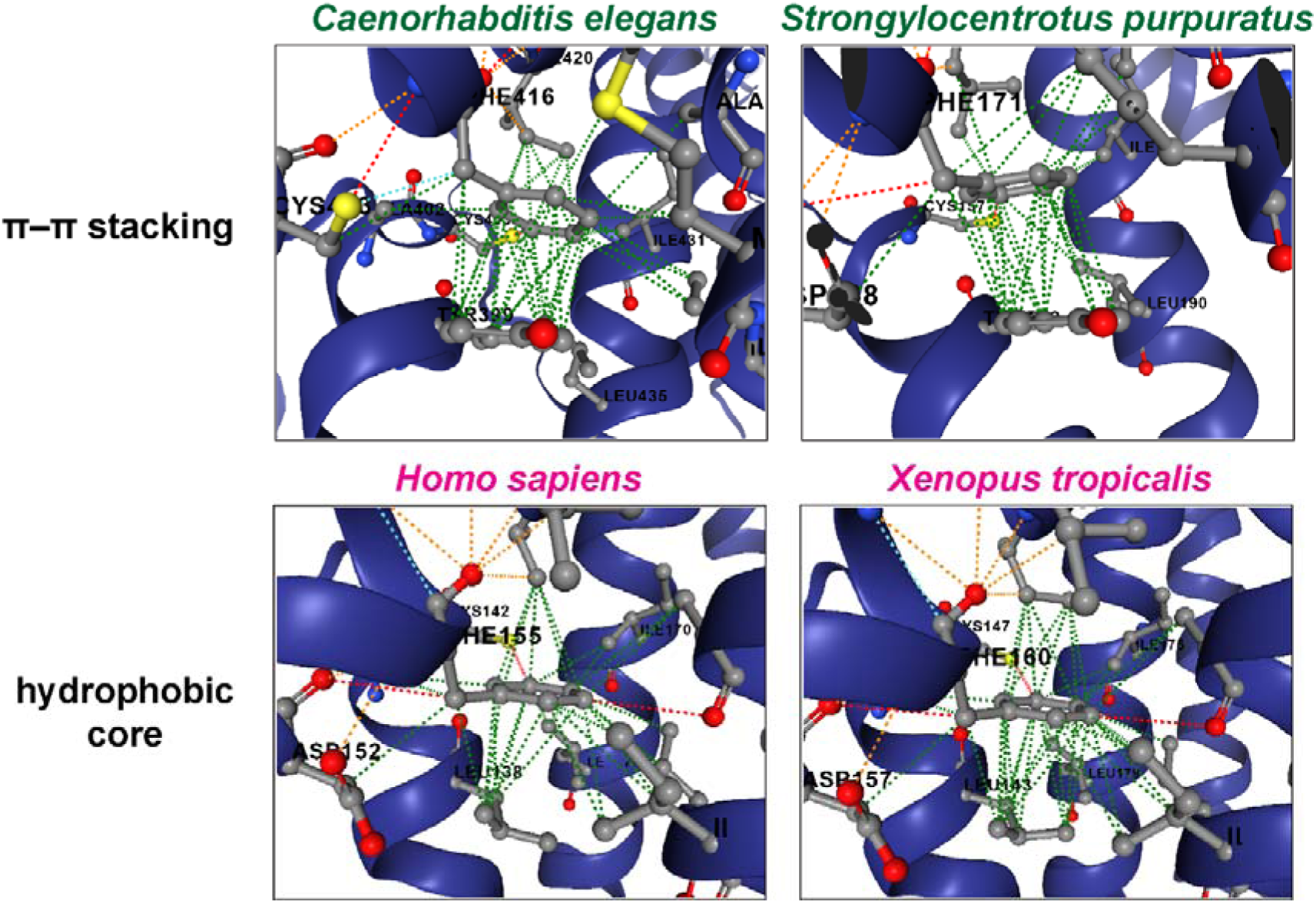
Conserved BAG domain architecture is maintained through distinct core stabilization mechanisms across evolution. In invertebrates (in green), including *C. elegans* and *S. purpuratus*, BAG domain stability is supported by π–π stacking interactions between aromatic residues (i.e., phenylalanine and tyrosine), forming a central stabilizing interaction within the domain core. In vertebrates (in pink), including humans and *X. tropicalis*, BAG domain stability is maintained through hydrophobic interactions between hydrophobic residues (i.e. phenylalanine and leucine), replacing aromatic stacking while preserving overall structural organization.

## Discussion

Our study integrates genetic, computational, and AlphaFold-based structural approaches to uncover how specific amino acids regulate UNC-23/BAG2 function in the context of muscle integrity and how these requirements have been conserved or altered through evolution. Consistent with the role of UNC-23 in muscle maintenance, worms carrying the C403Y mutation, which replaces a conserved cysteine, exhibit pronounced muscle deterioration. The conservation of cysteine 403 in UNC-23 (corresponding to cysteine 142 in human BAG2) across diverse animal phyla strongly suggests it has a fundamental role in the BAG domain’s function. In the context of BAG2’s co-chaperone activity, this cysteine is predicted to stabilize the interaction with Hsc70 (HSP-1 in worms), a major chaperone involved in protein folding and degradation. The UNC-23^C403Y^ mutation may impair UNC-23 function by altering structural or interaction properties of the BAG domain and, consequently, leading to muscle defects. Beyond this single conserved cysteine, our results dissected the structural architecture of the BAG domain to identify the amino acids most crucial for maintaining its three-dimensional fold. Using *in silico* stability predictions (DDMut analysis) and evolutionary conservation data, we pinpointed several hydrophobic residues in both UNC-23 and BAG2 that were predicted to be important for the protein’s core stability. Many of these residues (e.g., phenylalanine, leucine, isoleucine) are buried in the hydrophobic core, a common feature in proteins where packing of nonpolar side chains drives proper folding. Consistent with protein engineering studies, our analysis highlights that disrupting these core hydrophobic interactions tends to destabilize the protein structure. Indeed, phenylalanine 416 of UNC-23 (equivalent to phenylalanine 155 in human BAG2) emerged as a particularly critical core residue: replacing it with alanine (UNC-23^F416A^) caused a pronounced muscle deterioration phenotype *in vivo*, akin to a loss-of-function. This experimental result demonstrates that F416 is indispensable for UNC-23 function *in vivo* and consistent with a critical role in maintaining the BAG domain core.

Interestingly, while the UNC-23^F416A^ mutation had a drastic effect, the other three alanine substitutions targeting conserved core residues (UNC-23^C423A^, UNC-23^I431A^, UNC-23^L435A^) did not produce any obvious phenotypic defects under our experimental conditions. There are several possible reasons why these predicted stability-compromising mutations were apparently tolerated. First, the magnitude of destabilization caused by each substitution may not have been sufficient to impair UNC-23 function in the context of a living organism. Proteins are often marginally stable, and a modest decrease in stability might not translate into an immediate loss of function; cellular quality-control mechanisms or the presence of molecular chaperones could buffer the protein’s folding in these mutants. Second, it is possible that the functional assays we employed (focused on muscle integrity under standard laboratory conditions) were not sensitive to subtler defects. The UNC-23 mutants carrying C423A, I431A, or L435A might have mild biochemical or physiological impairments that do not manifest as obvious muscle deterioration, or they might become evident only under stress conditions or in aging animals. Another consideration is that not all highly conserved residues are critical for stability; some may be conserved for roles in interactions or regulation that were not assayed in our study. Our negative results for these three mutations, therefore highlight the importance of validating computational predictions experimentally. They demonstrate that conservation and modelling alone are not error-free indicators of *in vivo* importance and underscore the resilience and adaptability of protein structures, as the BAG domain can tolerate certain substitutions in evolutionarily conserved amino acids without collapsing.

One of the most illuminating findings from our comparative analysis is that the BAG domain’s AlphaFold-predicted overall fold is evolutionarily conserved from invertebrates to vertebrates, despite a fundamental shift in the nature of its core stabilizing interactions. In UNC-23 and other invertebrate BAG2 homologs, structural integrity appears to hinge on a π–π stacking interaction between two aromatic residues, e.g. phenylalanine 416 and tyrosine 399 in *C. elegans*. This interaction forms a rigid, centrally positioned aromatic stack that contributes significantly to core stabilization. In contrast, the vertebrate homologs, including human BAG2, use a phenylalanine–leucine pair (e.g. phenylalanine 155 and leucine 174 in human) at the analogous positions to stabilize the domain via purely hydrophobic packing. Our modelling (**Figure 7**) illustrates that this evolutionary substitution from a π–π stack to hydrophobic contact alters the chemical nature of the interaction but preserves the spatial and functional role within the domain’s hydrophobic core.

This shift reflects a broader evolutionary strategy observed across other protein families, wherein aromatic residue interactions, for instance π–π stacking or edge-to-face, are replaced by hydrophobic aliphatic side chains such as leucine or isoleucine, or vice versa, without compromising the protein’s global fold. For example SUMO–ubiquitin comparison demonstrate that a triad of aromatic residues in SUMO’s core, including a Tyr–Phe pair, can be functionally replaced in ubiquitin by a Leu–Ile–Leu hydrophobic cluster^33^. In this case, evolution maintains the packing density and hydrophobic character of the core, even while swapping chemically distinct side chains. Thus, our work suggests that π–π stacking and hydrophobic packing can serve as alternative stabilizing strategies for preserving BAG domain architecture during evolution.

This insight into divergent core interactions also sheds light on our mutational results. The critical importance of F416 in *C. elegans* UNC-23 can be rationalized by its unique role in the π–π stack with Y399. When F416 is replaced, this central stack is lost, profoundly destabilizing the domain. In contrast, residues like C423, I431, or L435, while conserved and contributory to the hydrophobic environment, do not participate in such a singular defining interaction in the worm protein, therefore, their individual loss may be buffered by the remaining structure. In the human BAG2 core, the phenylalanine–leucine interaction means that one would predict both F155 and L174 to be individually crucial, a prediction supported by our stability analysis showing BAG2^F155A^ and BAG2^L174A^ among the most destabilizing mutations. It will be interesting in the future to test whether mutating the human BAG2’s core phenylalanine (F155) has as dramatic an effect on BAG2 function as the UNC-23^F416A^ mutation did in worms.

In the broader context of protein structure–function relationships, our work underscores the value of integrating evolutionary and biophysical analyses to identify functionally critical residues. By combining computational predictions with *in vivo* experimental validation in the whole organism setting, we distinguished residues that are absolutely required for protein function from those that are less crucial, despite being conserved.

Finally, our findings have implications for disease research involving BAG2 and related co-chaperones. BAG2 is known to influence the progression of cancers (via stabilization of mutant p53)^14^ and to protect against neurodegenerative processes (by preventing tau protein aggregation)^19^. While no BAG2 mutations have yet been directly linked to human disease, our study in *C. elegans* UNC-23 suggests that mutations affecting the stability of the conserved BAG domain may likewise impair BAG2 function in humans. It is possible that subtle variants or post-translational modifications affecting BAG2’s core stability or its critical cysteine (C142 in humans) could impair its co-chaperone activity, potentially exacerbating proteostasis disorders. Moreover, the concept of distinct yet functionally equivalent core interactions in BAG2 raises the possibility that small-molecule stabilizers tailored to one mechanism (e.g. enhancing aromatic stacking or hydrophobic packing) could be designed to bolster BAG2 function if it were destabilized. In summary, by deciphering the amino acids crucial for UNC-23/BAG2 stability and function, we provide deeper insight into how this co-chaperone operates at a molecular level, advancing our fundamental understanding of BAG domain biology across evolution.

## Methods

### Worm Maintenance and Strains

*C. elegans* strains were cultured at 20□°C on nematode growth medium (NGM) plates seeded with *Escherichia coli* OP50 as a food source. Details of all strains used in this study are listed in Supplementary Table 2.

### EMS mutagenesis

EMS (ethyl methanesulfonate) mutagenesis was carried out essentially as described by Brenner^23^ with minor modifications. Synchronized L4 hermaphrodites were collected from well-fed NGM plates, washed three times in M9 buffer, and then incubated in 50 mM EMS diluted in M9 with gentle rotation at 20 °C for 4 h in a chemical fume hood. After exposure, worms were washed five times in M9 buffer until no trace of EMS remained, then transferred to fresh NGM plates seeded with *E. coli* OP50 to recover and lay eggs overnight. Approximately 100 P0 animals were used, and their F1 progeny were grown to maturity. Individual F1 hermaphrodites were singled onto separate plates to generate clonal F2 populations, which were subsequently screened for desired mutant phenotypes.

### Whole genome sequencing for mutation identification

#### Preparation of mutants and DNA extraction for WGS

The mutant strain was crossed with the original strain TUR59, which had been used for the EMS mutagenesis. 20 F2 recombinants exhibiting the same phenotype as the mutant strain were selected and transferred individually to separate NGM plates. Animals from 2-3 starved plates per each of the F2 lines were pooled, following the protocol described by Lehrbach et al^22^. Genomic DNA was then extracted using the Gentra Puregene Tissue Kit (Qiagen), according to the manufacturer’s instructions. DNA from two pools of DNA from two independent crosses was then used for whole genome sequencing.

#### Library preparation and whole genome sequencing (Novogene Inc.)

Genomic DNA was randomly sheared, end□repaired, A□tailed, and ligated to Illumina adapters. Fragments were size□selected, PCR□amplified, purified, quantified by Qubit and qPCR, and size distribution confirmed on a Fragment Analyzer. Libraries were pooled and sequenced on an Illumina NovaSeq X platform (150 bp PE, ∼3 Gb/sample). Raw reads were trimmed and filtered using fastp (v0.20.0) before alignment to the *C. elegans* WBcel235 reference genome (Ensembl 109) with BWA□MEM (v0.7.17). Sorted BAMs were merged, and variant calling was performed with GATK HaplotypeCaller (v4.0.5.1), followed by SNP/InDel filtering. Structural variants and copy□number variants were detected using BreakDancer (v1.4.4) and CNVnator (v0.3), respectively, and all variants were annotated with ANNOVAR. Candidate causative mutations were identified by comparing mutant and parental strain variant sets and prioritized based on predicted impact.

### Evolutionary Conservation and Phylogenetic Analysis

Multiple sequence alignment (MSA) of UNC-23/BAG2 homologs was performed using Clustal Omega^25^. Sequence visualization and phylogenetic analysis were conducted using Jalview version 2.11.4.1^34^. The following protein sequences were retrieved and utilized for the analysis: *Caenorhabditis elegans* CCD72352.1, *Homo Sapiens* CAG38527.1, *Mus musculus* NP_663367.1, *Bos taurus* NP_001029436.1, *Monodelphis domestica* XP_056670323.1, *Ornithorhynchus anatinus* XP_001505275.1, *Gallus gallus* XP_001505275.1, *Danio rerio* NP_957294.2, *Xenopus tropicalis* AAH75476.1, *Apis mellifera* XP_623942.2, *Strongylocentrotus purpuratus* XP_795015.2.

### Generation of *unc-23* mutant strains

*Unc-23* point mutants were generated by microinjection of pre⍰assembled Cas9 RNP complexes and single⍰stranded repair templates, essentially as described by Dokshin et al.^24^. Briefly, gene⍰specific CRISPR RNA (crRNA) and trans⍰activating crRNA (tracrRNA) (IDT) were mixed with recombinant Cas9 nuclease (IDT Alt⍰R S.p. Cas9) and incubated at 37 °C for 10 min to form ribonucleoprotein (RNP) complexes. A 100–200 nt single⍰stranded oligodeoxynucleotide (ssODN) template bearing the desired nucleotide substitution and silent PAM⍰blocking mutations was added directly to the RNP mix, together with 50 ng/µL pRF4□rol⍰6(su1006)] as a co⍰injection marker. Young adult hermaphrodites were injected using the standard technique^35^. Injected P0 animals were recovered to OP50⍰seeded NGM plates at 20°C. Roller F1 progeny were singled, and the target locus was PCR□amplified and Sanger□sequenced to identify correctly edited alleles; homozygous F2 lines were confirmed by sequencing.

The sequences of all oligonucleotides used in this study are listed in Supplementary Table 3.

### Developmental timing assay

To assess developmental rates, populations of wild type (ACH93), UNC-23^C403Y^ (TUR262), and UNC-23^E297K^ mutants (TUR274) strains were synchronized by hand-picking embryos at the late 3-fold stage of embryonic development. Synchronized individuals were cultured on standard Nematode Growth Medium plates seeded with *E. coli* OP50 and incubated at 20°C for 72 hours. Following incubation, worms were imaged using a Zeiss Axio Zoom.V16 stereomicroscope equipped with an Axiocam 705 Mono camera at 70x magnification. Based on the acquired images, the developmental stage of each individual was determined, and the proportion of animals in L4 larval versus adult stages was quantified for each strain.

### Morphometric analysis and muscle width quantification

For morphometric quantification, animals were age-synchronized by hand-picking L4 larvae and incubating them for 24 hours at 20°C to reach the first day of adulthood. Day 1 adults were imaged at 70× magnification using a Zeiss Axio Zoom.V16 stereomicroscope equipped with an Axiocam 705 Mono camera. All measurements were performed using Fiji software (ImageJ version 2.16.0/1.54p)^36^. Quantitative analysis included the width of both the dorsal and ventral body wall muscles, using the position of the uterus as a landmark to consistently identify the ventral side and ensure anatomical uniformity across all samples. Whole head width was measured at the level of the pharyngeal isthmus.

### Computational and Structural Analysis

#### Protein Structural Models

Three-dimensional protein models were retrieved from the AlphaFold Protein Structure Database^37^. The specific entries used for analysis were AF-O61980-F1-model_v4 (*Caenorhabditis elegans*), AF-O95816-F1-model_v4 (*Homo sapiens*), AF-A0A7M7HFP3-F1-model_v4 (*Strongylocentrotus purpuratus)*, and AF-Q3ZBG5-F1-model_v4 (*Bos taurus*). *Structural alignment and RMSD calculation*

Global and local structural alignments were performed using the Matchmaker tool in UCSF ChimeraX 1.9^38,39^. Alignments were calculated based on the “Best-aligning pair” of chains using the Needleman-Wunsch algorithm and the BLOSUM-62 similarity matrix. To assess the conservation of the core architecture, an initial root-mean-square deviation (RMSD) was calculated across all 197 atom pairs. This was refined by performing a secondary alignment on a pruned subset of 99 residues corresponding to the BAG domain and its immediate flanking sequences to calculate the final RMSD, effectively excluding high-variance regions such as flexible termini.

#### Interaction visualization and analysis

Visualization and characterization of internal protein interactions were conducted using DDMut^30^. π–π stacking interactions in invertebrates were identified based on the facial orientation between the aromatic rings of phenylalanine and tyrosine residues. Hydrophobic core interactions were assessed by identifying clusters of non-polar side chains (e.g., phenylalanine, leucine, and isoleucine) and visualizing the loss of contacts upon alanine substitution.

### *In silico* protein stability predictions

The impact of single-point mutations on the thermodynamic stability (ΔΔG) of the UNC-23/BAG2 was predicted using DDMut, a deep learning-based tool designed to estimate stability changes upon mutation^30^. For these calculations, protein structural models were retrieved from the AlphaFold Protein Structure Database^37^. The DDMut server was utilized to perform a systematic alanine scan, where every residue within the UNC-23 and BAG2 proteins was computationally substituted with alanine to assess its contribution to structural stability. The resulting predictions were used to rank the substitutions by their destabilizing potential, focusing on the top 15 most destabilizing residues (those with the highest predicted decrease in stability) for subsequent analysis^30^.

### Microscopic imaging

Representative fluorescence images were acquired using a Leica STELLARIS 8 FALCON confocal system with a 40× oil immersion objective. GFP and RFP fluorescence were excited using 494 nm and 575 nm lasers, respectively. Worms were immobilized on 5% agarose pads for imaging, using either 10□μl of 0.05□μm polystyrene microspheres (PolySciences) or 25□μM tetramisole. Representative bright-field images were captured at 70x magnification on a Zeiss Axio Zoom.V16 stereomicroscope equipped with an Axiocam 705 Mono camera, using ZEN software for image acquisition.

### Statistical Analysis

No prior statistical methods were applied to determine sample sizes. Statistical significance was assessed using the Kruskal-Wallis test with Dunn’s multiple comparisons test and Fisher’s exact test. A p-value of < 0.05 was considered statistically significant.

## Acknowledgements

Some strains were provided by the CGC, which is funded by NIH Office of Research Infrastructure Programs (P40 OD010440). We thank Filip Kozłowski and Tomasz Strumiński for assistance with worm maintenance. Work in the MT Laboratory was mainly funded by a National Science Centre SONATA grant (2019/35/D/NZ3/04091 to MT) and additionally supported by a National Science Centre SONATA BIS grant (2021/42/E/NZ3/00358 to MT).

## Author Contributions

Conceptualization: MT; Data curation: MT; Formal analysis: MŚ, MT, RPS, AS; Funding acquisition: MT; Investigation: MŚ, RPS, AS, MT; Methodology: MT; Project administration: MT; Resources: MT; Supervision: MT; Validation: MT; Visualization: RPS, MŚ, AS, MT; Writing – original draft: MT; Writing – review & editing: MT, RPS, AS, MŚ.

